# The Development of Wet Laboratory Methods for Improved Downstream Analysis of High Throughput Sequencing Data for Virus and Viroid Diagnostics of *Vitis vinifera* in Australian Post-Entry Quarantine

**DOI:** 10.1101/2025.02.26.640443

**Authors:** Stephanie I. Morgan, Emmanuelle C. McFarlane, Naima Tasnim, Bhuwaneshwariba Vala, Dilani DeSilva, Ruvini V. Lelwala

## Abstract

High throughput sequencing (HTS) methods are an essential tool in Australia’s Post-Entry Quarantine (PEQ) facility to facilitate the rapid, reliable and sensitive diagnostics of plant viruses and viroids. Small RNA sequencing (sRNAseq) for the detection of viruses and viroids has been implemented for several plant commodities in PEQ, including clonal grasses, *Prunus* spp., *Rubus* spp., and *Fragaria* spp., with the potential to expand to further plant genera. Currently, imported grapevine (*Vitis* spp.) require many PCR assays (22) which are both laborious and time consuming. In this study we compare different wet laboratory processes for *Vitis vinifera* ribonucleic acid (RNA) extraction, including different tissue types, tissue weights and elution volumes and their effects on detecting viruses and viroids by Illumina sRNA sequencing. Overall, our results suggest that increased RNA concentration of *V. vinifera* can be achieved by using cambium tissue, reducing elution volume and decreasing the tissue weight. Furthermore, when comparing sRNAseq to current PCR methods in post-entry quarantine, sRNAseq outperformed the prescribed PCRs.

## 1. Introduction

Grapevine (*Vitis* spp.) is one of the most economically significant plant species worldwide [1]. The continual importation of new *Vitis* spp. varieties is crucial for Australian grapevine industries to remain competitive and profitable, by allowing them to be adaptive to meet consumer demands and changing climate [2]. However, the importation of new *Vitis* spp. varieties from overseas is also a major pathway for the entry of pathogens of biosecurity significance into Australia [3]. Viruses and viroids of *Vitis* spp. can cause significant economic losses for Australian agricultural industries, including the wine industry [4]. These viruses and viroids can be detrimental for *Vitis* spp., reducing yield, fruit quality and plant vigour [4]. For example, the spread of *Grapevine Pinot gris virus* (GPGV) in Europe has been associated with reduced yield and quality of grapes, as well as symptoms impacting fruit and plant growth such as delayed budburst, stunted shoots, restricted spring growth, zig-zag shoots and millerandage [5-11]. While GPGV was first detected in Australia in 2016 there is yet no clear correlation between GPGV presence and disease symptoms demonstrated to date and the full impact of this virus on Australian nursery and fruit production is still somewhat unknown [5]. Similarly, *Grapevine red blotch virus* (GRBV), which causes significant economic losses in the USA (US$2,231 to US$68,548 per hectare) due to its impact on berry quality [12], was first reported in Australia in 2022 and its impact to Australia still unknown [13]. Due to the similarity of Australia’s climate to other grapevine growing regions worldwide, and the significance of grapevine viruses in other parts of the world, GPGV, GRBV and many like them have significance to Australia’s grapevine industries [14].

Rapid and robust diagnostic methods are an essential tool in facilitating the importation of new *Vitis* spp. germplasm into Australia, by preventing the introduction of exotic plant pathogens of biosecurity significance. To sustain demand, there is a need for diagnostics to evolve and become more effective and efficient. The current Australian post entry quarantine (PEQ) requirements for *Vitis* spp. include pathogen screening using 22 molecular polymerase chain reaction (PCR) assays [14]. Moreover, pathogens of biosecurity significance, and in particular viruses, are continuously reviewed and new PCR targets are added every few years with the emergence of novel pathogens. The number of PCR assays is becoming increasingly difficult to sustain, particularly as *Vitis* spp. are known to be infected with more than 60 distinct virus species and this number continues to grow with the increased confidence in novel detections through high throughput sequencing (HTS) technologies [15]. HTS methods are increasingly replacing time-intensive, and specific PCR methods and provide additional benefits to traditional diagnostic methods [16]. Due to the large volume of data generated, HTS methods have the potential to detect low titre pathogens, related pathogens, as well as novel pathogens, traits that are highly desirable for futureproofing biosecurity screening [17].

To date, HTS methods have not been applied to all plant genera in Australian PEQ. This is partly due to the stringent validation and verification requirements prior to implementation, demonstrating that HTS standards for sensitivity, reproducibility and specificity can be met [18]. Additionally, HTS methods must prove cost effective when compared to more traditional diagnostic methods such as PCR and Enzyme-Linked Immuno-Sorbent Assay (ELISA). With HTS becoming more affordable, this requirement is in turn becoming more achievable [18]. HTS has become increasingly used for the diagnostics of plant viruses and viroids [18]. Small RNA sequencing (sRNAseq) has been successfully implemented in Australian PEQ, enabling viroid and virus screening in a single assay [18] for *Prunus* spp., *Fragaria* spp., *Rubus* spp. and clonal grasses [14]. Moreover, sRNAseq has facilitated the identification of novel viruses [19], and is better suited to detect new viral strains compared to traditional methods, allowing for detection of pathogens despite mutations [15]. These additional benefits of sRNAseq enable Australian biosecurity systems to be better equipped for future plant pathogen threats.

The success of sRNAseq heavily relies on achieving good RNA quality, integrity and yield. This has proven challenging for woody plants, including *Vitis* spp., compared to herbaceous plants. This is largely due to the presence of secondary metabolites, including polysaccharides and polyphenols in woody plants [19]. These secondary metabolites react with RNAs, forming insoluble complexes and remaining as contaminants in the nucleic extract. This reaction results in reducing the RNA quality and yield of these woody plants and thus impacting downstream test methods, such as sRNAseq [1,19].

The objective of this study was to compare different wet-laboratory processes to achieve improved integrity and quantity of total RNA from *V. vinifera*, suitable for performing sRNAseq. Moreover, the variations tested had to be compatible with the current methods used at the Australian PEQ to seamlessly be integrated in existing extraction workflows, due to the large diversity of plant commodities routinely tested at the Australian PEQ. A series of experiments were conducted to compare (A) tissue types, (B) tissue weights, and (C) elution volumes, on RNA concentration and integrity using the Promega Maxwell robotic system. Additionally, multiple extracts from two virus positive grapevine plants were sequenced using sRNAseq and viral detections were compared between sRNAseq and PCR diagnostic methods.

## Materials and Methods

### 2.1. Sample Preparation

Two *V. vinifera* cultivars, Variety 1 (V1) and Variety 2 (V2) growing at the Australian Post-Entry Quarantine facility, Mickleham, were selected for wet laboratory optimisation after prescribed testing via PCR revealed the presence of viruses. The plants were grown in a BC2 glasshouse with a set temperature of 26 °C and 50-85% humidity. Both leaf and cambium were sampled from three locations on the plant (top, middle and lower branches) to ensure a representative sample was obtained. Material was sampled and kept chilled at 4 °C for transportation to the laboratory and kept on ice during sub-sampling.

Cambium and midrib were dissected from the chilled tissue using single-use sterile scalpel blades and subsamples (20 mg and 50 mg) were weighed on a Sartorius Quintix 1102-1S precision balance and collected into 2 mL Eppendorf safe-lock tubes containing two 7 mm stainless steel beads and stored at –80 °C for a minimum of 30 minutes. The samples were collected in replicates as outlined in Supplementary Table 2.

### 2.2. RNA Extraction

RNA extraction was performed using the Promega Maxwell RSC Simply RNA Tissue Kit (AS1430) and the Promega Maxwell RSC instrument as per the manufacturer’s protocols [20], except for the following deviations. Tissuelyser LT 12-tube adaptor racks were kept at –80 °C for a minimum of 30 minutes prior to homogenisation of sub-sampled *V. vinifera* tissue. Chilled Homogenisation Solution (200 µL) was added to each tube containing the previously frozen *V. vinifera* tissue before disrupting the tissue for 1 minute using the Tissuelyser LT (oscillation frequency 50 Hz). The tubes were briefly pulsed in a chilled microcentrifuge and a further 400 µL of chilled Homogenisation Solution was added to each sample before processing for another minute in the Tissuelyser LT (50 Hz). Lysis Buffer (300 µL) was added to each tube, vortexed and then centrifuged for 5 minutes at 20,000 G in a chilled microcentrifuge. Supernatant (500 µL) from each sample tube was added to the allocated cartridge in well number one as per manufacturer’s instructions. Additionally, a synthetic RNA spike-in, cel-miR-39 (3.3 fmol/mL) (Norgen Biotek) was added into well number 1 as an internal control for the extraction of sRNA and downstream HTS sequencing analysis. Different volumes (50 µL and 100 µL) of double-distilled water were used for final elution of RNA and the volumes of cel-miR39 solution added to well 1 was adjusted to 2 µL and 4 µL respectively.

### 2.3. RNA quality assessment and quantification

RNA quality scores (RINe - RNA Integrity Number equivalent) and concentration of RNA samples were assessed using an Agilent 4200 TapeStation System to evaluate RNA extraction optimisation success prior to sending extracts to the external sequencing service provider. Sample preparation and instrument operation were conducted as per the manufacturer’s protocols [21].

Twelve extracts (including an additional leaf/ midrib sample) were carefully selected for sRNAseq, based on their concentrations and RINe scores, to increase success during the library preparation stage for sRNAseq (Sup. Table 2). Total RNA extracts (17 µL) were shipped on dry ice to the external service provider.

### 2.4. Library Preparation and sRNA Sequencing

RNA samples were processed by an external service provider according to the optimised protocols reported in previous work [18].

Briefly, the HTS service provider repeated quality control checks using either a NanoDrop or Epoch Spectrophotometer to measure the absorbance of nucleic acids and determine the potential contaminants prior to library preparation. The HTS service provider recommends that samples should meet RNA integrity scores greater than 8 and concentration greater than 20 ng/µL.

Samples that did not meet the required RINe and concentration requirements were concentrated by vacuum to meet library preparation input of 100 ng per sample. QIAseq miRNA 96 Index IL UDI-A kit with unique dual indexes (UDIs) and unique molecular IDs (UMI) was used for sRNA library preparation (Qiagen). QIAseq FastSelect-rRNA plant kit (Qiagen) was used during library preparation to remove abundant host plant RNA. Sequencing was performed on an Illumina Novaseq X plus instrument for 75 cycles aiming to generate a minimum of 40 million raw reads and 4 million sRNA reads per sample. Demultiplexed FASTQ files provided by the HTS service provider were used for downstream bioinformatics analysis at the PEQ facility.

### 2.5. Bioinformatics Analysis

sRNAseq data was analysed using VirReport and GA-VirReport bioinformatics workflows published previously [18, 23]. Fragments Per Kilobase of transcript, per Million mapped reads (FPKM) was used to compare pathogen titre as FPKM is the normalised unit of transcript expression and scales by transcript length to compensate for RNA-Seq protocols often generating more sequencing reads from longer RNA molecules, that were not of interest in this experiment.

### 2.6. Statistical Analysis

An additional TapeStation dataset (concentration and RINe) derived from 29 *V. vinifera* plants extracted previously during prescribed testing was combined with that of V1 and V2 extracted in this study, and statistical analysis performed on all 31 plants (Supp. Table 3). There was a total of 61 extracts analysed with some plants extracted singly, whilst others were extracted with replication (Supp. Table 3).

All statistical analyses were performed in Genstat using an unbalanced ANOVA (Analysis of Variance) on the entire TapeStation dataset, to account for unequal sample sizes. Homogeneity of variance was determined by examining plots of fitted values versus residuals, while histograms of residuals were examined for normality of distribution (Supp. Fig. 1). The level of significance used was P ≤ 0.05.

## 2. Results

### 3.1. Effect of wet laboratory optimisation on RNA Integrity Number equivalent (RINe)

As RINe scores were only generated for samples with a concentration above 10 ng/µL, and thus only generated for 20/61 sample extracts, this data was excluded from further statistical analysis (Supp. Table 3). Even in samples with low concentrations, when TapeStation electropherograms were examined, there were two distinct peaks for 18S and 28S, (Fig. 1).

**Figure 1.**
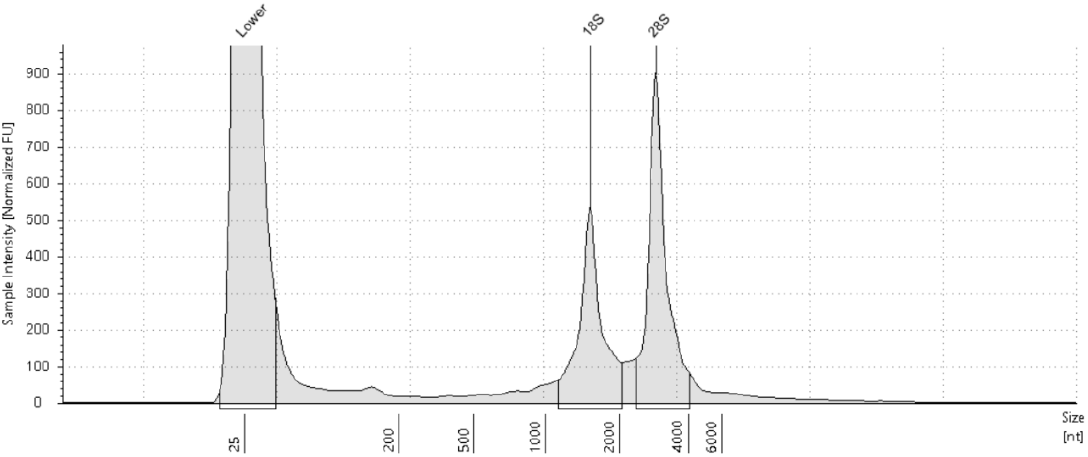
Electropherogram of an extract from V1 with low concentration (9.43 ng/µL) that was sent for sRNAseq. Electropherogram is scaled to sample. 28S peak is approximately double the size of the 18S peak and there are no other obvious peaks, showing good RNA integrity.

### 3.2. Effect of wet laboratory optimisation on RNA concentration

For the TapeStation dataset, the interaction between tissue type, tissue weight and elution volume were not significant (P=0.101), regarding yield (RNA concentrations). Moreover, the interaction between tissue type and elution volume (P=0.922), tissue type and tissue weight (P=0.837), and elution volume and tissue weight (P=0.153) were also not significant regarding yield.

#### 3.2.1. Tissue type on RNA concentration

RNA extracted from cambium tissue had a greater average concentration of RNA than RNA extracted from midrib, 12.83±8.47 ng/µL and 2.26±0.14 ng/µL for V1, and 7.02±4.19 ng/µL and 3.51±1.28 ng/µL for V2, respectively (Fig. 2, Supp. Table 2). When comparing tissue types, midrib and cambium, the same relationship was observed across both *V. vinifera* varieties (Fig. 2, Supp. Table 2).

**Figure 2.**
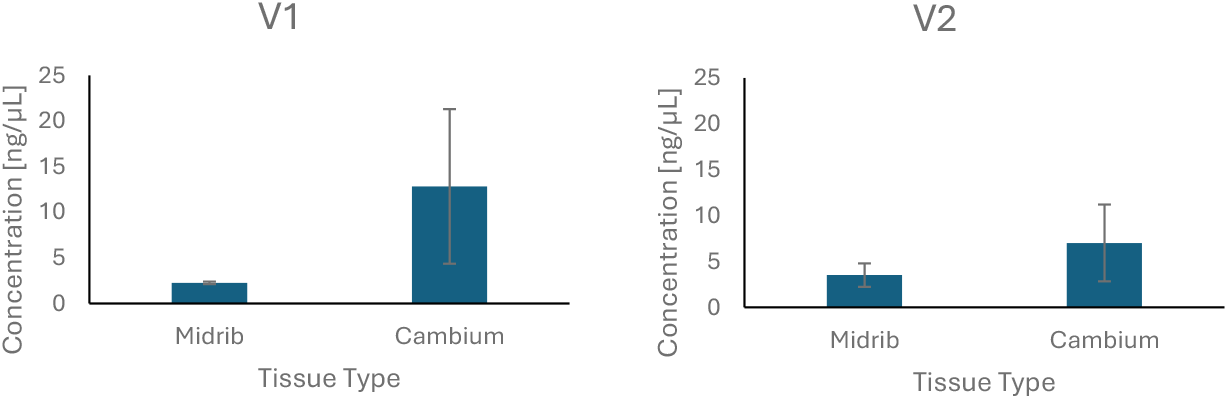
The effect of tissue type on RNA concentration for V1 and V2. Blue histograms represent the average concentration for each tissue type and standard deviation bars are displayed.

The TapeStation dataset from 31 *V. vinifera* showed there was a significant difference between tissue type (P<0.001), with cambium having a higher concentration than midrib (Fig. 3).

**Figure 3.**
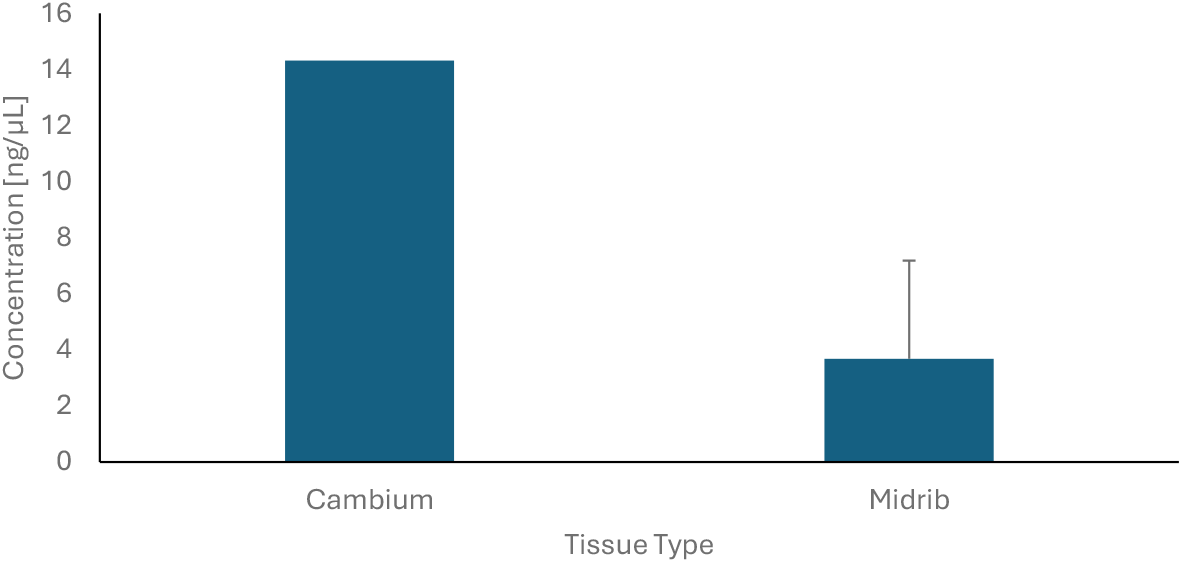
The effect of tissue type on RNA concentration for 31 *V. vinifera* plants. Blue histograms represent median concentration for each tissue type and Least Significant Difference (LSD) bars are displayed above (P=0.05).

#### 3.2.2. Elution volume on RNA concentration

RNA eluted in a volume of 50 µL had an average concentration greater than RNA eluted in 100 µL, 17.45±10.02 ng/µL and 6.52±3.70 ng/µL for V1, and 6.76±4.53 ng/µL and 4.47±2.59 ng/µL for V2, respectively (Fig. 4, Supp. Table 2). There was a negative relationship between elution volume and RNA concentration in both V1 and V2 (Fig. 4, Supp. Table 2).

**Figure 4.**
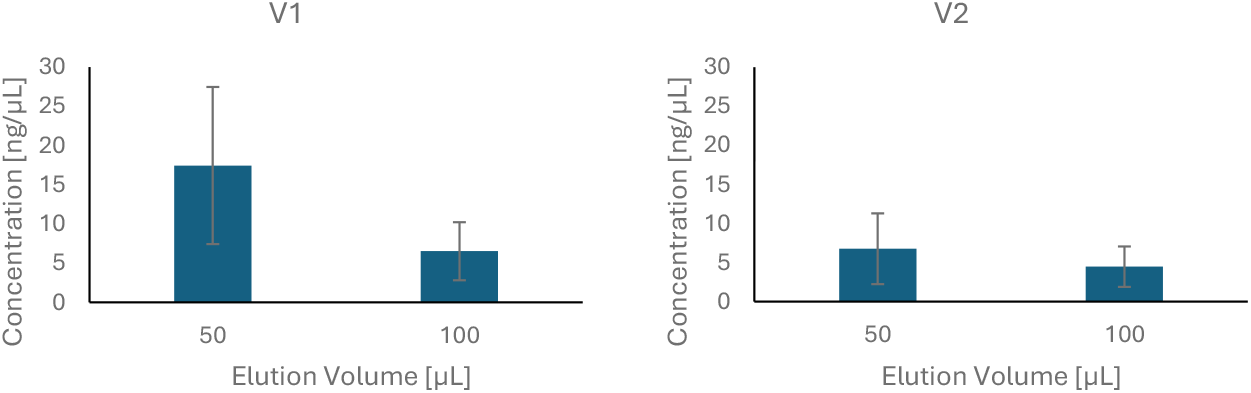
The effect of elution volume on RNA concentration for V1 and V2. Blue histograms represent the average concentration for each elution volume and standard deviation bars are displayed.

When the TapeStation dataset from 31 *V. vinifera* was analysed statistically, the concentration was significantly higher with an elution volume of 50 µL (P<0.001, Fig. 5).

**Figure 5.**
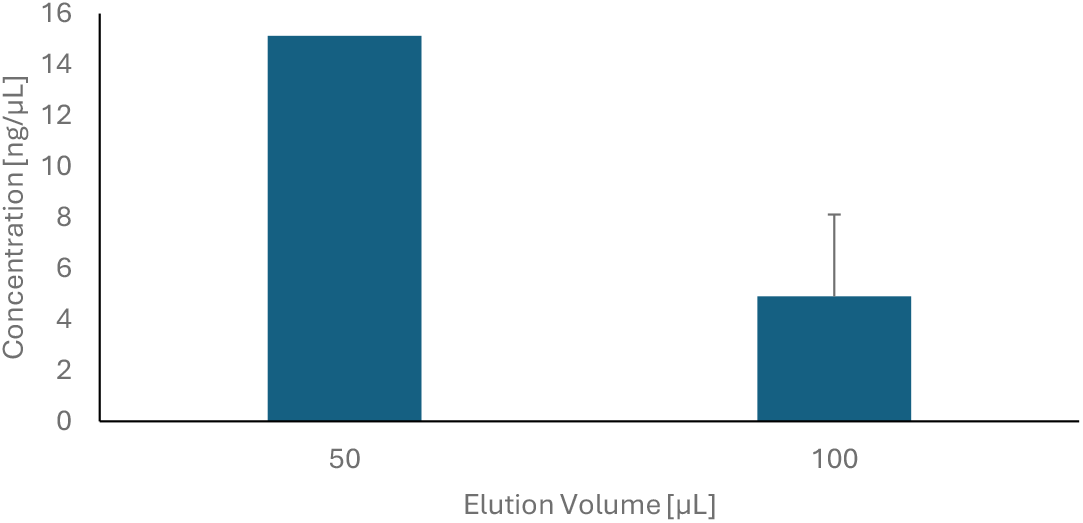
The effect of elution volume on RNA concentration for 31 *V. vinifera* plants. Blue histograms represent median concentration for each elution volume and LSD bars are displayed above (P=0.05).

#### 3.2.3. Tissue weight on RNA concentration

RNA extracted from a tissue weight of 20 mg had a greater average concentration than RNA extracted from a tissue weight of 50 mg, 14.81±11.01 ng/µL and 7.33±3.43 ng/µL for V1, and 6.72±4.56 ng/µL and 4.50±2.58 ng/µL for V2, respectively (Fig. 6, Supp. Table 2). There was a negative correlation between tissue weight and concentration (Fig. 6, Supp. Table 2).

**Figure 6.**
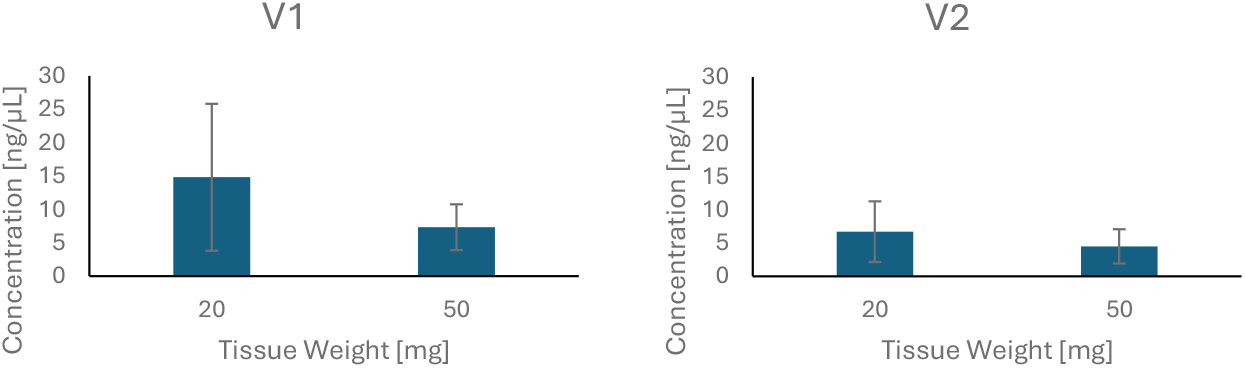
The effect of tissue weight on concentration for *V. vinifera* V1 and V2. Blue histograms represent the average concentration for each tissue weight and standard deviation bars are displayed.

The TapeStation dataset, comprised of 31 *V. vinifera* plants, showed that the yield extracted from samples with a tissue weight of 20 mg was significantly greater than the yield extracted from samples with a tissue weight of 50 mg (P=0.005).

### 3.3. Effect of wet laboratory optimisation on sRNAseq parameters

Despite the suboptimal RNA concentrations, all 12 RNA extracts resulted in successful miRNA libraries. All samples except a single replicate of the V1 – cambium generated more than 43 million reads per sample. All samples, including the one with 37.7 million raw reads retained more than 4.48 million informative reads after quality control (QC) processing, filtering for plant reads and filtering for 21-22 nt length. PEQ currently uses a threshold of minimum 4 million informative reads per sample, to be adequate for viral detections. Therefore, all samples were determined sequenced to an adequate depth for viral diagnostics.

The NanoDrop data generated by the HTS sequencing provider had on average better A_260/280_ ratios and A_260/230_ ratios for cambium tissue than midrib tissue for both V1 and V2 (Supp. Table 4.). On average, V1 had a ratio of absorbance at A260 and 280 nm (the A_260/280_ ratio) of 2.07±0.06 for cambium and 1.45±0.07 for midrib. V2 had on average a A_260/280_ ratio of 1.75±0.35 for cambium and 1.53±0.19 for midrib. The average A_260/230_ ratios for V1 in cambium were 1.40±0.10 and in midrib 0.65±0.07. The average A_260/230_ ratios for V2 in cambium were 0.90±0.28 and in midrib 0.65±0.13.

sRNAseq detected eight additional pathogens in V1 than prescribed PCR testing (Table 1), including targets not tested through PCR and in one instance, a pathogen target that was not amplified by PCR, *Grapevine berry inner necrosis virus* (Table 1). Similarly, sRNAseq detected six pathogens in V2 additional to those detected by the PCR assays prescribed in Australian post-entry quarantine (Table 1). Detections were consistent across both tissue types (midrib and cambium) in both V1 and V2.

**Table 1.**
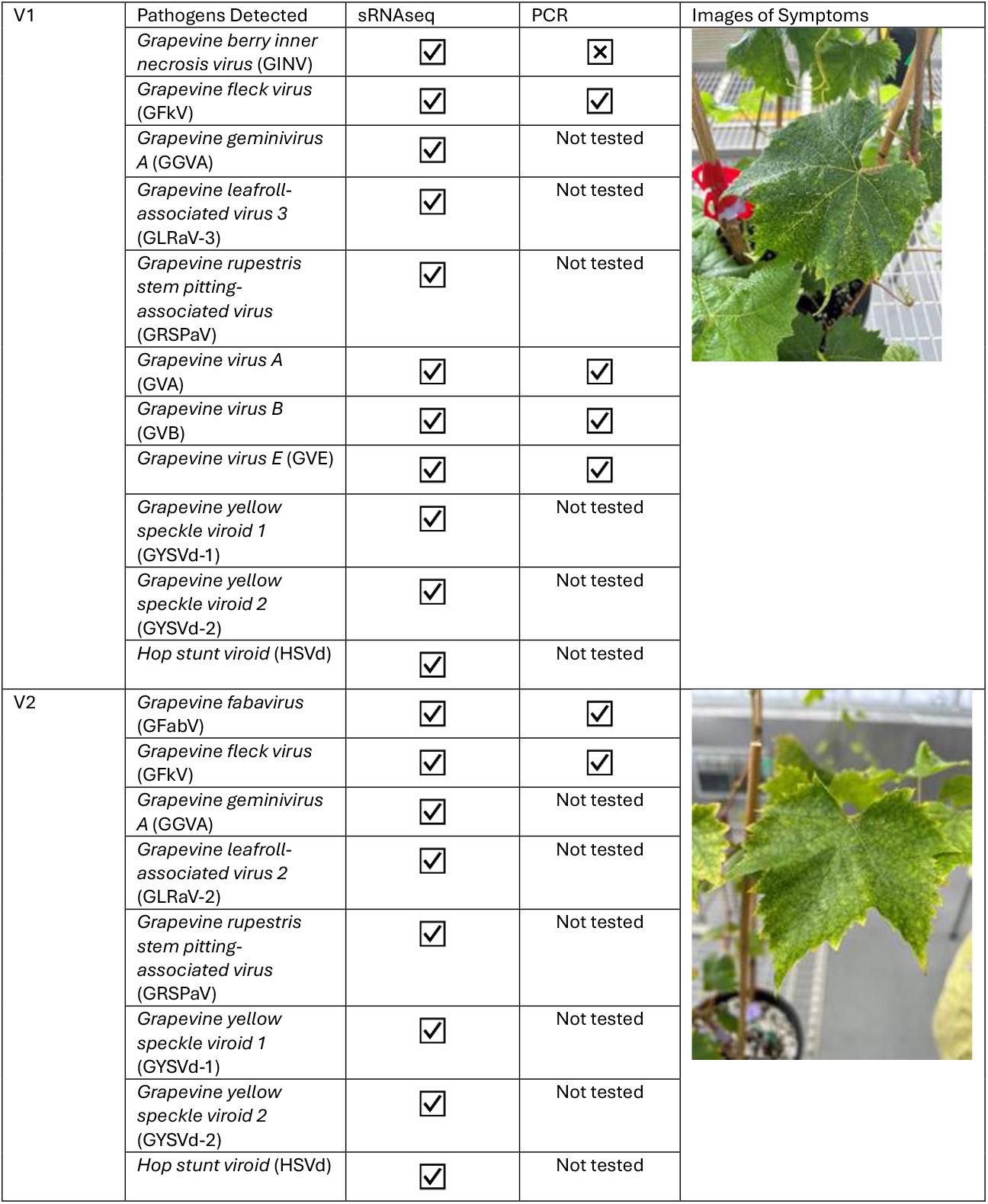
Pathogens detected in V1 and V2 using sRNAseq and PCR, and images of symptomatic material. A tick indicates that the pathogen was detected using the method listed, a cross indicates that the pathogen was not detected using the method listed.

| V1 | Pathogens Detected | sRNAseq | PCR | Images of Symptoms |
| --- | --- | --- | --- | --- |
|  | *Grapevine berry inner necrosis virus* (GINV) | ✓ | ✗ |  |
|  | *Grapevine fleck virus* (GFKV) | ✓ | ✓ |  |
|  | *Grapevine geminivirus A* (GGVA) | ✓ | Not tested |  |
|  | *Grapevine leafroll-associated virus 3* (GLRaV-3) | ✓ | Not tested |  |
|  | *Grapevine rupestris stem pitting-associated virus* (GRSPaV) | ✓ | Not tested |  |
|  | *Grapevine virus A* (GVA) | ✓ | ✓ |  |
|  | *Grapevine virus B* (GVB) | ✓ | ✓ |  |
|  | *Grapevine virus E* (GVE) | ✓ | ✓ |  |
|  | *Grapevine yellow speckle viroid 1* (GYSVd-1) | ✓ | Not tested |  |
|  | *Grapevine yellow speckle viroid 2* (GYSVd-2) | ✓ | Not tested |  |
|  | *Hop stunt viroid* (HSVd) | ✓ | Not tested |  |
| V2 | *Grapevine fabavirus* (GFabV) | ✓ | ✓ |  |
|  | *Grapevine fleck virus* (GFKV) | ✓ | ✓ |  |
|  | *Grapevine geminivirus A* (GGVA) | ✓ | Not tested |  |
|  | *Grapevine leafroll-associated virus 2* (GLRaV-2) | ✓ | Not tested |  |
|  | *Grapevine rupestris stem pitting-associated virus* (GRSPaV) | ✓ | Not tested |  |
|  | *Grapevine yellow speckle viroid 1* (GYSVd-1) | ✓ | Not tested |  |
|  | *Grapevine yellow speckle viroid 2* (GYSVd-2) | ✓ | Not tested |  |
|  | *Hop stunt viroid* (HSVd) | ✓ | Not tested |  |

When looking at the summarised detections generated from the two HTS pipelines (VirReport and GA-VirReport), cambium tissue generated on average a greater FPKM for eight out of 11 detections in V1, and for five out of eight detections in V2 (Fig. 6 and 7, Supp. Table 5).

**Figure 7.**
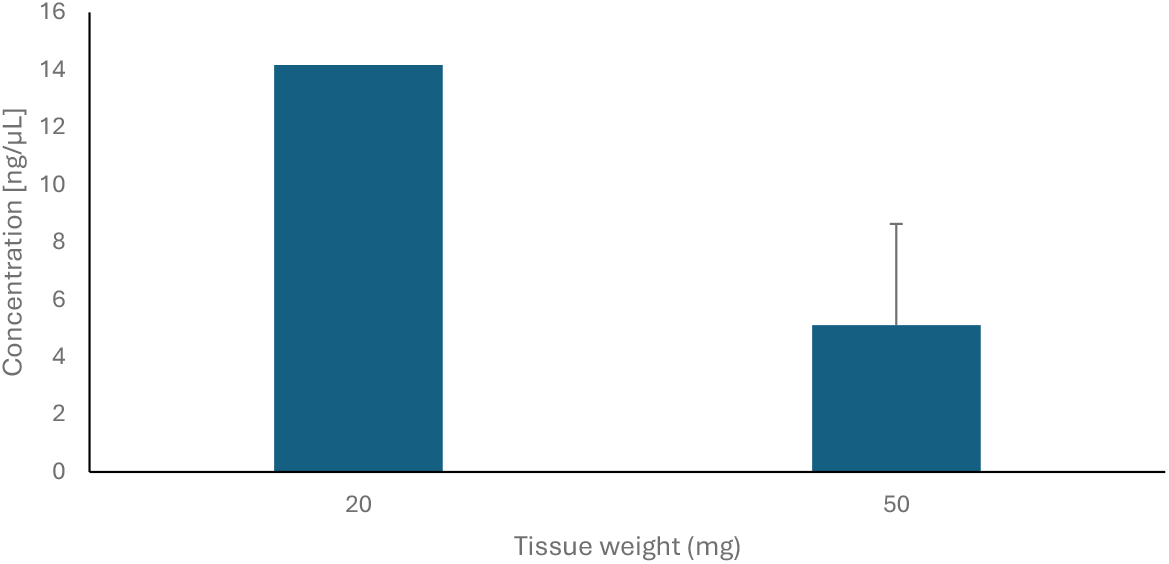
The effect of tissue weight on concentration for 31 *V. vinifera* plants. Blue histograms represent median concentration for each tissue weight and LSD bars are displayed above (P=0.05).

**Figure 6.**
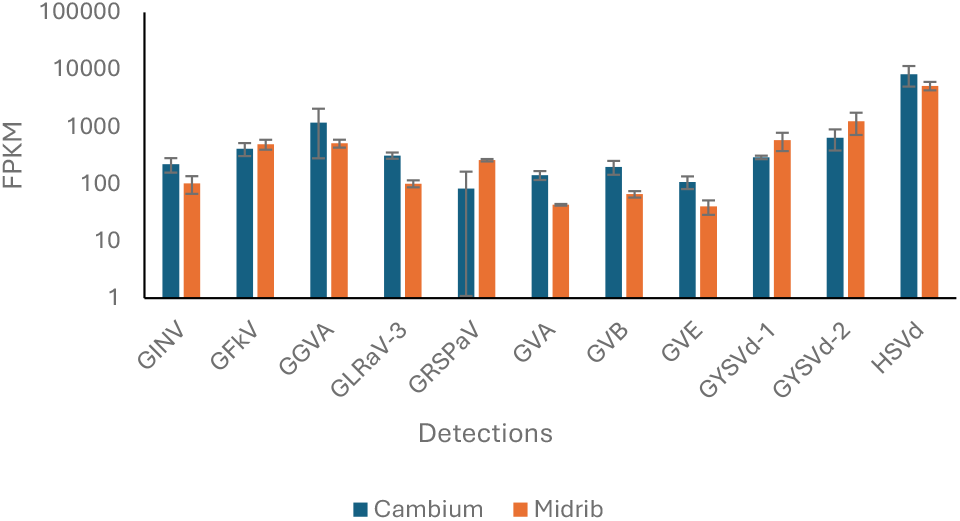
sRNAseq detections comparing cambium and midrib tissue types for V1. FPKM values are from the VirReport workflow and listed on a logarithmic scale. Standard deviation bars are displayed.

**Figure 7.**
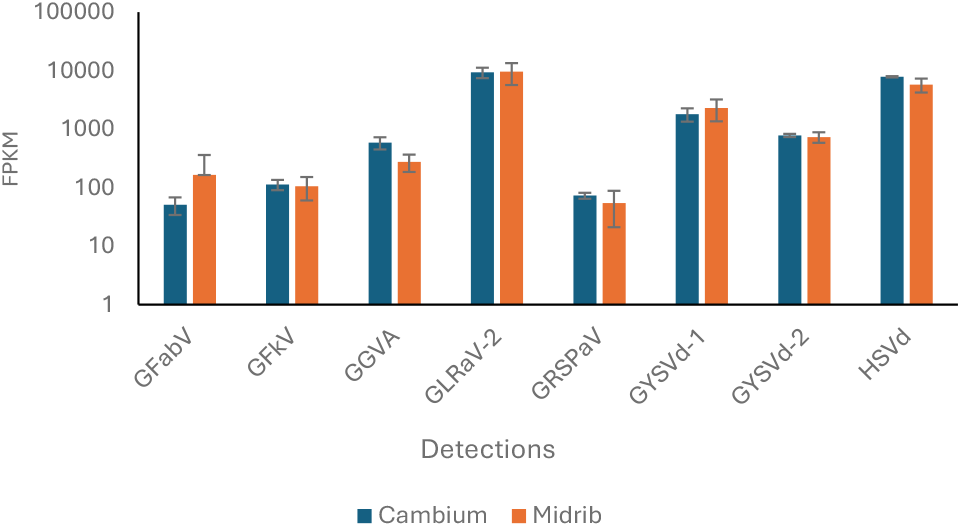
HTS detections comparing cambium and midrib tissue types for V2. FPKM values are from the VirReport workflow and listed on a logarithmic scale. Standard deviation bars are displayed.

## Discussion

This study investigated changes in tissue type, tissue weight and elution volume to achieve better total RNA extractions of *V. vinifera* using the Maxwell robotic system, aiming to obtain RNA of sufficient quality and concentration for downstream sRNAseq detection of viral pathogens. The results of this study showed a significant positive correlation between cambium and RNA concentration, a negative correlation between elution volumes tested and RNA concentration, and a negative correlation between tissue weights tested and RNA concentration. These inferences were apparent for both V1 and V2, and for the TapeStation dataset.

Achieving good quality and yield RNA in *V. vinifera* is difficult due to the presence of high polysaccharides and phenolic compounds in the plant tissues. Phenolic compounds oxidise and bind to nucleic acids, causing polyphenols to be leftover in the extraction process, and subsequently, resulting in decreased RNA yield [22]. Several techniques, often laborious and time intensive, have been trialled over the years to increase quality of *Vitis spp*. nucleic acid extractions, with low to moderate success. Most techniques are only appropriate for low number of samples (6-12) and require several hours to several days to complete [24, 25, 26], making them ill-suited to the large volume of plant samples processed at the Australian PEQ facility.

Low quality and yield RNA was observed in this study, especially in *V. vinifera* RNA extracted from leaf tissue using a tissue weight of 50 mg and a final elution volume of 100 µL which is the currently used standard at PEQ. The NanoDrop data for cambium had an average A_260/280_ ratio of 1.91, which fell within the ideal range of 1.8-2, compared to midrib, which had an average A_260/280_ ratio of 1.49. As the A_260/280_ ratio is an indication of the RNA purity, a lower value could indicate that more proteins, contaminants or residual phenols are present in the midrib tissue than in cambium tissue [27]. Grapevine canes, which can be directly compared to the cambium tissue used in this study, have been shown to consistently produce better quality DNA extracts than leaves when several DNA extraction techniques were tested [24], likely because canes have been shown to have a lower secondary metabolite content and higher level of antioxidants than grapevine leaves [28]. Antioxidants play an important role in reducing the binding of nucleic acid to oxidised polyphenols. To this effect, polyvynilpyrrolidone (PVP) was also incorporated into the RNA extraction method in the current study to prevent the oxidation of phenolic compounds and secondary metabolites during the extraction, in turn preventing the resulting tannins and melanins from covalently bonding to nucleic acid and precipitating in the final elute [22, 24]. Cambium tissue also produced on average greater pathogen titre than midrib tissue, further showing its suitability in downstream viral diagnostics through sRNAseq.

Reducing the elution volume is a practice that is often used to concentrate nucleic acid extractions. Indeed, there is an obvious negative correlation between concentration and volume, with a reduced volume resulting in increased concentration. Through previous optimisation work using the Promega Maxwell system, and due to the large number of tests required for PEQ diagnostics, the standard elution volume used at the Australian PEQ is 100 µL. However, as the elution volume had a significant impact on RNA concentration for *V. vinifera* in this study, the effects of reducing the elution volume were assessed and found acceptable. As the current concentration requirements for QC checks and sRNAseq are 20 µL, using an elution volume of 50 µL provides more than double the requirement for diagnostic testing.

Interestingly, the smallest weight of tissue tested resulted in a significantly higher concentration in this study. Similarly to what was observed with the differences in tissue type, it is likely that a smaller weight would have resulted in less secondary metabolites such as polysaccharides and phenolic compounds, thus increasing nucleic acid yield. For PEQ diagnostics, leaf material is generally preferred as it does not impact the growth and health of young plants as much as cambium sampling [29]. Requiring a smaller quantity of tissue would enable a smaller sample to be taken from the plant and perhaps facilitate the sampling of cambium tissue for PEQ diagnostics. Additionally, if cambium samples are taken during pruning activities, no harm would be done to the plant. Implementing these changes in diagnostics would aid in the adoption of sRNAseq in *V. vinifera* plants in PEQ.

In this study, more viral pathogens were detected through sRNAseq than the existing prescribed PCR testing, despite suboptimal QC scores in some instances. This could be explained by the fact that unlike PCR and qPCR, the HTS approach currently used at PEQ is non-specific and unbiased, and can therefore, be utilised as a screening tool for all viruses and viroids [30]. Moreover, HTS can be more sensitive than qPCR [30]. This was evident for *Grapevine berry inner necrosis virus*, which was detected only through sRNAseq, but not through current PCR testing methods, noting that the current protocol uses midrib tissue and not cambium tissue. The detection of *Grapevine berry inner necrosis virus* through sRNAseq led to further optimisation of the current assay to detect the virus, and retesting of the grapevine plants through the optimised assay. This demonstrates the advantage sRNAseq provides in screening for pathogens, rather than targeted PCR assays that may not allow for the detection of new strains and novel pathogens. Furthermore, this added benefit of sRNAseq positions Australian biosecurity to be better equipped for tackling new and emerging biosecurity threats.

A consideration for future work would be to increase the number of replicates and include plants of known pathogen status, to determine whether the entire virome is able to be detected. Additionally, by increasing the number of plants in the study, the implications of high viral load on efficiency, if any, may be inferred. A future objective is to establish the minimum quality control thresholds for *Vitis* spp. for sRNAseq diagnostics.

## 4. Conclusions

Overall, sampling cambium tissue, sampling 20 mg tissue weight and using 50 µL elution volume each provided increased total RNA concentrations. Sampling cambium tissue also improved RNA quality. sRNAseq outperformed PCR, by detecting more virus and viroid pathogens, and providing quicker processing times, thus proving to be a more effective diagnostic tool. The outcomes of this study will influence the future of *V. vinifera* virus and viroid diagnostics in Australian post-entry quarantine, and its effectiveness in protecting Australian *Vitis* spp. from harmful pathogens.

## Author contributions

Conceptualisation, S.I.M., E.C.M., and R.V.L.; methodology, S.I.M., E.C.M., and R.V.L.; writing – original draft preparation, S.I.M.; writing – review and editing, S.I.M., E.C.M., R.V.L., N.T., B.V., and D.D. All authors have read and agreed to the published manuscript.

## Acknowledgements

The authors acknowledge the horticulture team at the Australian PEQ for growing and maintaining the plants that were sampled for this study. We would also like to acknowledge and thank Dr. Dylan McFarlane, Research Scientist, VSICA Research, for assisting in the statistical analysis of our data.

