## Supplemental Data for "The Development of Wet Laboratory Methods for Improved Downstream Analysis of High Throughput Sequencing Data for Virus and Viroid Diagnostics of *Vitis vinifera* in Australian Post-Entry Quarantine"

Supplementary Data

**Supplementary Table 1.** Molecular PCR assays for virus and viroid prescribed *V. vinifera* diagnostics in Australian PEQ.

| Pathogen Target | Primer Sequence (5'-3') | Reference |
|---|---|---|
| *Grapevine fabavirus* | F – ATTTTGCAGCAAGCICCATGGG<br>R – TCCAAAGACCAGAACTAAACCTC | Chiaki Y., *et al.* (2020) |
| *Tobacco necrosis virus A*<br>*Tobacco necrosis virus D* | F – AAGAYWCAAYACATTWCDATCG<br>R – TGGTTCAATGAACTCAATCWCGTA | Xi, D., *et al.* (2008) |
| *Grapevine angular mosaic virus*<br>*Grapevine line pattern virus* | F – GCNGGWTGYGGDAARWCNAC<br>R – AMDGGWAYYTGYTYNGTRTCACC | Univertos, M., *et al.* (2010) |
| *Grapevine deformation virus*<br>*Grapevine fanleaf virus*<br>*Raspberry ringspot virus – grapevine strain* | F – AATAAATCATAA-ACDTCWGARGGITAYCC<br>R – RATDCCYACYTGRCWIGGCA | Wei, T., *et al.* (2008). |
| *Artichoke Italian latent virus*<br>*Grapevine Anatolian ringspot virus*<br>*Grapevine chrome mosaic virus*<br>*Tomato black ring virus* | F – AATAAATCATAATCTGGITTTGCYTTRACRGT<br>R – CTTRTCACTVCCATCRGTAA | Wei, T., *et al.* (2008). |
| *Blueberry leaf mottle virus New York strain*<br>*Grapevine Bulgarian latent virus*<br>*Grapevine Tunisian ringspot virus*<br>*Peach rosette mosaic virus*<br>*Tomato ringspot virus* | F – TTRKDYTGGYKAAMYYCCA<br>R – TMATCSWASCRHGTGSKKGCCA | Digiaro M., et al. (2007) |
| *Petunia asteroid mosaic virus* | F – AAGGGTAAGGATGGTGAGGA<br>R - TTTGGTAGGTTGTGGAGTGC | Harris, R., *et al.* (2007) |
| *Grapevine berry inner necrosis virus* | F – TTGTCTGATGGCTCTGATG<br>R - ACRGTKGATARACASGTGG | Cho, I.S., et al. (2013) |
| *Grapevine asteroid mosaic associated virus* | F – CYCARCAYAARGTVAACGA<br>R - GCGCATGCABGTSAGRGGG | Beaino, T., *et al.* (2001) |
| *Grapevine virus B*<br>*Grapevine virus D*<br>*Grapevine virus E*<br>*Grapevine virus G*<br>*Grapevine virus H*<br>*Grapevine virus I*<br>*Grapevine virus J*<br>*Grapevine virus K*<br>*Grapevine virus L*<br>*Grapevine virus M* | F – GGNGARTTYGGNACNTTYTTYTTY<br>R - NCKCCANCCRCARAANARNGG | Diaz-Lara A., *et al.* (2010) |
| *Cherry leafroll virus (CLRV) – grape isolate* | F – TGGCGACCGTGTAACGG<br>R1 – TACTACTAAGACCGGTCGCATGG<br>R2 – TACTACTAAGACCGGTCGCATGAA<br>P1 – FAM- GTTAAGGTGACACTGGTGG -MGB NFQ<br>P2 – FAM- TTACGGTGACACTGGTGG-MGB NFQ | Osman, F., *et al.* (2014) |
| *Citrus yellow vein clearing virus* | F – TCCAACTCACAAACCCAGTG<br>R – ATGGGCTC TTGGTTTTCCTT | Hongming C., *et al* (2016) |
| *Grapevine leafroll associated virus-7* | F – TATATCCCAACGGAGATGGC<br>R – ATGTTCCTCCACCAAAATCG | Engel EA., *et al.* (2008) |
| *Grapevine leafroll associated virus-13* | F – CGAAGGTTTGCTATACGTCAACCC<br>R - TGAACAAGTGACGCCGACGA | Ito, T., *et al.* (2016) |
| *Grapevine red blotch virus* | F – AAGAAGCCGCAGGAGTC<br>R – CCAGACGACGTCTTTGAA<br>P - [6FAM]-TTCACGTGCAGCATTTAATATT-[BHQ1] | Agriculture Victoria (AgriBio) unpublished |
| *Grapevine Roditis leaf discolouration-associated virus* | F1 - AGTTTCTTCAACAAGTCAGC<br>F2 – AGTTTCTTCAACAAATCAGC<br>F3 – AGCTTCTTCAACAAATCCG<br>R1 – TGATTGCTCRTTTAACTGG<br>R2 – GACTGTTCCTTCAGCTG | Morán F., *et al.* (2020) |

| | | |
|---|---|---|
| | P - 6-FAM-<br>ATTAACAGTTTCTTTCTTACAGAATGGAA-BHQ1-<br>ZNA4 | |
| *Grapevine vein clearing virus* | F – GTAAACCTCATGACTCTCATG<br>R – CTTCTCCTTCAGAAATTGAGCAGAT | Guo Q., *et al.*<br>(2014) |
| *Grapevine virus T* | F – ATGTAYTACTCYAARGTRATATGG<br>R - ATTGTARGCTGGRGCDCCC | Diaz-Lara A., *et al.*<br>(2020) |
| *Strawberry latent ringspot sadwavirus* | F – GCTCTTTGCTTTCTTTTGTGTT<br>R – TCTAAGTGCCAGAACTAAACC<br>P – FAM- CTGGGAGGATGCCTGGTTAATCCTTT -<br>31ABkFQ | Ministry for<br>Primary Industries<br>(2016)<br>unpublished |

**Supplementary Table 2.** Raw Tapestation values and replication information for *V. vinifera* Varieties 1 and 2.

| Tissue Type | Variety (1,2) | Tissue Weight [mg] | Elution Volume [μL] | Concentration [ng/μL] | RINe | 28S/18S (Area) | Sequenced by sRNAseq (Y/N) |
|---|---|---|---|---|---|---|---|
| Midrib | 2 | 20 | 50 | 4.48 | | | Y |
| Midrib | 2 | 20 | 100 | 3.87 | | | Y |
| Midrib | 1 | 20 | 100 | 2.36 | | | Y |
| Cambium | 1 | 20 | 50 | 31.2 | 8.9 | 1.7 | N |
| Cambium | 1 | 20 | 100 | 4.6 | | | N |
| Cambium | 1 | 20 | 50 | 23.2 | 8.6 | 1.5 | Y |
| Cambium | 1 | 20 | 50 | 12.1 | 9 | 1.3 | Y |
| Cambium | 1 | 20 | 50 | 15.4 | 8.9 | 1.2 | Y |
| Cambium | 2 | 20 | 50 | 13.8 | 8.1 | 1.4 | Y |
| Cambium | 2 | 20 | 50 | 8.73 | | | Y |
| Cambium | 2 | 20 | 50 | 2.74 | | | N |
| Midrib | 2 | 50 | 50 | 4.05 | | | Y |
| Midrib | 2 | 50 | 100 | 1.63 | | | Y |
| Midrib | 1 | 50 | 100 | 2.16 | | | Y |
| Cambium | 1 | 50 | 50 | 5.34 | | | N |
| Cambium | 1 | 50 | 100 | 9.43 | | | N |
| Cambium | 1 | 50 | 100 | 5.93 | | | N |
| Cambium | 1 | 50 | 100 | 11.2 | 8.9 | 1 | N |
| Cambium | 1 | 50 | 100 | 9.93 | | | N |
| Cambium | 2 | 50 | 100 | 3.65 | | | N |
| Cambium | 2 | 50 | 100 | 8.68 | | | N |
| Cambium | 2 | 50 | 100 | 4.51 | | | N |

**Supplementary Table 3.** Raw values and replication information for TapeStation dataset of *V. vinifera* used in statistical analyses. Plant/cultivar 1 represents Variety 1 and Plant/cultivar 2 represents Variety 2. Variables in blue were tested for their effects on categories in pink.

| Plant/cultivar | Tissue type | Tissue weight (mg) | Elution volume (µL) | RINe score | Concentration (ng/ µL) |
|---|---|---|---|---|---|
| 1 | cambium | 20 | 50 | 8.9 | 31.2 |
| 1 | cambium | 20 | 100 | | 4.6 |
| 1 | cambium | 50 | 50 | | 5.34 |
| 1 | cambium | 50 | 100 | | 9.43 |
| 1 | cambium | 20 | 50 | 8.6 | 23.2 |
| 1 | cambium | 20 | 50 | 9 | 12.1 |
| 1 | cambium | 20 | 50 | 8.9 | 15.4 |
| 1 | cambium | 50 | 100 | | 5.93 |
| 1 | cambium | 50 | 100 | 8.9 | 11.2 |
| 1 | cambium | 50 | 100 | | 9.93 |
| 1 | midrib | 20 | 100 | | 2.36 |
| 1 | midrib | 50 | 100 | | 2.16 |
| 2 | cambium | 20 | 50 | 8.1 | 13.8 |
| 2 | cambium | 20 | 50 | | 8.73 |
| 2 | cambium | 20 | 50 | | 2.74 |
| 2 | cambium | 50 | 100 | | 3.65 |
| 2 | cambium | 50 | 100 | | 8.68 |
| 2 | cambium | 50 | 100 | | 4.51 |
| 2 | midrib | 20 | 50 | | 4.48 |
| 2 | midrib | 20 | 100 | | 3.87 |
| 2 | midrib | 50 | 50 | | 4.05 |
| 2 | midrib | 50 | 100 | | 1.63 |
| 2 | midrib | 20 | 100 | | 2.33 |
| 3 | midrib | 50 | 100 | | 2.14 |
| 4 | midrib | 50 | 100 | | 1.9 |
| 5 | midrib | 50 | 100 | | 1.98 |
| 6 | midrib | 50 | 100 | | 2.23 |
| 7 | midrib | 50 | 100 | 7.2 | 10.7 |
| 8 | midrib | 50 | 100 | | 2.62 |
| 9 | midrib | 50 | 100 | | 2.61 |
| 10 | midrib | 50 | 100 | | 2.55 |
| 11 | midrib | 50 | 100 | 7.7 | 20.3 |
| 12 | midrib | 50 | 100 | | 5.41 |
| 13 | midrib | 50 | 100 | | 1.71 |
| 14 | midrib | 50 | 100 | | 2.91 |
| 15 | midrib | 50 | 100 | | 5.15 |
| 16 | midrib | 50 | 100 | | 1.64 |
| 17 | midrib | 50 | 100 | | 2.63 |

| | | | | | |
|---|---|---|---|---|---|
| 18 | cambium | 20 | 50 | 7.9 | 24.4 |
| 18 | midrib | 50 | 100 | | 2.36 |
| 19 | cambium | 20 | 50 | 8.8 | 14.8 |
| 19 | midrib | 50 | 100 | | 2.79 |
| 20 | cambium | 20 | 50 | | 4.8 |
| 20 | midrib | 50 | 100 | | 4.63 |
| 21 | cambium | 20 | 50 | 8.9 | 29.1 |
| 21 | midrib | 50 | 100 | | 7.39 |
| 22 | cambium | 20 | 50 | | 8.25 |
| 22 | cambium | 20 | 50 | | 8.48 |
| 22 | cambium | 20 | 50 | | 4.89 |
| 22 | midrib | 20 | 50 | | 2.56 |
| 22 | midrib | 20 | 50 | | 1.97 |
| 22 | midrib | 20 | 50 | | 1.92 |
| 23 | cambium | 20 | 50 | 8.2 | 22.8 |
| 24 | cambium | 20 | 50 | 8.5 | 19.9 |
| 25 | cambium | 20 | 50 | 8.2 | 27.9 |
| 26 | cambium | 20 | 50 | 8.8 | 29.2 |
| 27 | cambium | 20 | 50 | 8.7 | 26.2 |
| 28 | cambium | 20 | 50 | 8.5 | 28 |
| 29 | cambium | 20 | 50 | 8 | 25.4 |
| 30 | cambium | 20 | 50 | 8.4 | 40 |
| 31 | cambium | 20 | 50 | 8.4 | 17.1 |

**Supplementary Figure 1.** Graphs of residuals used to assess the suitability of an unbalanced ANOVA for statistical analysis of TapeStation dataset.

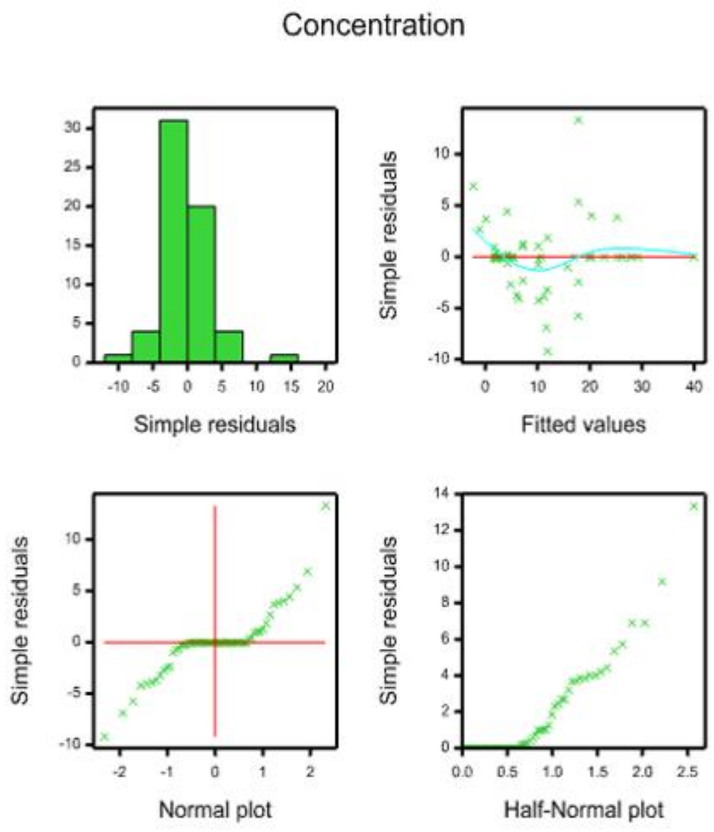


**Supplementary Table 4.** Quality data for the *V. vinifera* sRNAseq samples, performed by the sequencing provider.

| Variety | Tissue Type | Concentration Epoch/Nanodrop (ng/µL) | 260:280 Epoch | 260:230 Epoch |
|---|---|---|---|---|
| V2 | Midrib | 4 | 1.4 | 0.5 |
| V2 | Midrib | 4 | 1.4 | 0.7 |
| V1 | Midrib | 3 | 1.4 | 0.7 |
| V1 | Cambium | 30 | 2.0 | 1.5 |
| V1 | Cambium | 19 | 2.1 | 1.4 |
| V1 | Cambium | 18 | 2.1 | 1.3 |
| V2 | Cambium | 24 | 2.0 | 1.1 |
| V2 | Cambium | 16 | 1.5 | 0.7 |
| V2 | Midrib | 8 | 1.8 | 0.8 |
| V2 | Midrib | 2 | 1.5 | 0.6 |
| V1 | Midrib | 3 | 1.5 | 0.6 |

**Supplementary Table 5.** Average FPKM values for pathogen detections from the VirReport workflow for Varieties 1 and 2.

| Variety | Detections | Cambium (FPKM) | Standard deviation (cambium) | Midrib (FPKM) | Standard deviation (midrib) |
|---|---|---|---|---|---|
| V1 | *Grapevine berry inner necrosis virus* (GINV) | 219.33 | 62.07 | 101.50 | 34.65 |
| | *Grapevine fleck virus* (GFkV) | 411.00 | 105.26 | 493.00 | 97.58 |
| | *Grapevine geminivirus A* (GGVA) | 1172.00 | 892.32 | 511.00 | 80.61 |
| | *Grapevine leafroll-associated virus 3* (GLRaV-3) | 313.67 | 39.31 | 101.00 | 14.14 |
| | *Grapevine rupestris stem pitting-associated virus* (GRSPaV) | 82.33 | 81.24 | 259.00 | 12.73 |
| | *Grapevine virus A* (GVA) | 142.00 | 25.24 | 43.00 | 1.41 |
| | *Grapevine virus B* (GVB) | 198.00 | 53.78 | 66.00 | 8.49 |
| | *Grapevine virus E* (GVE) | 108.00 | 26.96 | 40.00 | 11.31 |
| | *Grapevine yellow speckle viroid 1* (GYSVd-1) | 290.50 | 20.51 | 578.00 | 203.65 |
| | *Grapevine yellow speckle viroid 2* (GYSVd-2) | 640.00 | 257.12 | 1236.50 | 519.72 |
| | *Hop stunt viroid* (HSVd) | 8281.33 | 3245.09 | 5197.50 | 874.69 |
| V2 | *Grapevine fabavirus* (GFabV) | 51.00 | 16.97 | 164.20 | 196.98 |
| | *Grapevine fleck virus* (GFkV) | 113.00 | 22.63 | 106.00 | 45.66 |
| | *Grapevine geminivirus A* (GGVA) | 587.00 | 138.59 | 275.80 | 90.61 |
| | *Grapevine leafroll-associated virus 2* (GLRaV-2) | 9318.50 | 1883.03 | 9549.20 | 3878.38 |
| | *Grapevine rupestris stem pitting-associated virus* (GRSPaV) | 73.00 | 8.49 | 54.60 | 33.58 |
| | *Grapevine yellow speckle viroid 1* (GYSVd-1) | 1803.00 | 461.03 | 2286.20 | 928.41 |
| | *Grapevine yellow speckle viroid 2* (GYSVd-2) | 780.00 | 49.50 | 732.80 | 149.65 |
| | *Hop stunt viroid* (HSVd) | 7846.50 | 195.87 | 5761.00 | 1541.31 |
